# Undruggable oncoproteins cMyc and NMyc bind to mediator of transcription with superior high affinity

**DOI:** 10.1101/2024.03.15.585150

**Authors:** Andrea Knight, Josef Houser, Tomas Otasevic, Vilem Juran, Vaclav Vybihal, Martin Smrcka, Martin Piskacek

## Abstract

The overexpression of MYC genes is frequently found in many human cancers including adult and pediatric malignant brain tumors. Targeting MYC genes continues to be challenging due to their undruggable nature. The nine-amino-acid activation domain (9aaTAD) has been identified using our prediction algorithm in all four Yamanaka factors including c-Myc and showed to activate transcription as short peptides. We generated a set of c-Myc constructs (1-108, 69-108 and 98-108) in the N-terminal regions and tested their ability to initiate transcription. We discovered strong interactions in nanomolar scale between the 9aaTAD of c-Myc and N-Myc proteins with the KIX domain of CBP coactivator. The c-Myc 9aaTAD (region 98-108) was not overlapping with the MBII (region 128-143) and therefore represents TRRAP independent activation region. Next, we showed the 9aaTADs in human c-Myc and N-Myc conservation within the MYC family. Interestingly, the loss of the 9aaTAD in L-Myc paralogs was identified in higher metazoans suggesting the deletions had occurred in early tetrapod evolution. In summary, as c-Myc is largely intrinsically disordered protein and therefore difficult to target by small molecule inhibitors, our finding of the c-Myc 9aaTAD in complex with the KIX domain represents a promising druggable target for development of new peptide inhibitors in MYC-driven tumors.

## Introduction

The MYC (also known as c-Myc) oncogene, located on chromosome 8q24.21, encodes a transcription factor that normally regulates the expression of approximately 15% of human genes implicated in proliferation, growth, differentiation, metabolism, and stemness [1–3]. Myc is one of the four Yamanaka factors (c-Myc, Oct4, Sox2, and Klf4) that could reprogram differentiated somatic cells into pluripotent stem cells. Aberrant c-Myc expression has been observed in ∼70% of human cancers by driving autonomous cell cycle progression, recruitment of inflammatory cells, blocking differentiation, extensive stromal remodeling, invasion, and angiogenesis. Thus, MYC is thought to be a central driver of initiation of tumorigenesis [4–7].

Myc regulates broad transcriptional networks that controls developmental and homeostatic processes through direct activation of gene transcription, chromosomal translocation, genomic amplification, mRNA upregulation, and retroviral integration [8–11]. Structurally, MYC contains a basic helix-loop-helix leucine-zipper (bHLH-LZ) motif, mediating dimerization with its obligate partner, the Myc-associated protein x (MAX), and DNA binding [12–14].

The c-Myc protein harbors conserved regions MB0 (10-32), MBI (44-63), MBII (128-143), MBIIIa (188-199), MBIIIb (259-270), MBIV (304-324) and C-terminal DNA binding domain bHLHZip [15–17]. The molecular dynamic simulation for c-Myc (protein conformation combined with NMR experiments), provided the first insights into intrinsically disordered N-terminal region including MB0, MBI, MBII and activation domain regions [17]. The MBII region is responsible for interaction with general transcriptional coactivator TRRAP [18], which is component of human SAGA complex [19]. Noteworthy, EZH2 protein, a subunit of Polycomb repressive complex[2, is essential for oncogenesis of MLL rearranged leukaemias. The c-Myc (central region 144-349) build a complex with EZH2 and mediated activation of non-canonical EZH2 targets [20].

Despite four decades of research and drug development, c-Myc has been recognized as highly undruggable oncotarget [21,22]. Apart from the DNA binding domain, the c-Myc protein is largely unstructured and intrinsically disordered protein [23], and is linked with difficulty to design small molecule inhibitors. One of the most extensively investigated MYC inhibitors is a peptide OmoMYC, which acts by blocking of the c-Myc binding [15,24,25]. The OmoMYC peptide (OMO-103) is in the first-in-human phase I/II clinical trial in patients with advanced solid tumors (VHIO-born spin-off Peptomyc S.L., Spanish Agency of Medicines and Medical Devices, EU-funded project SYST-iMYC), (https://ClinicalTrials.gov/show/NCT04808362).

Recent study shows that the coactivator EZH2 promotes c-Myc-driven oncogenesis by binding to the c-Myc protein. Both EZH2 and c-Myc are efficiently destroyed by the small drug MS177, which targets the EZH2 protein [20]. It was discovered that the small drugs JQ1 and dBET6, which target the BET4 protein, also inhibit MYC expression [26–28]. Mitotic Aurora kinase A (AURKA) functions as a transcription factor stabilizes and binds c-Myc and N-Myc [29,30].

Previously, we had identified the nine-amino-acid activation domains (9aaTAD) using our prediction algorithm (www.med.muni.cz/9aaTAD) in numerous transcription factors including members of the Gal4-p53-E2A-MLL-SREBP-NF-SP-KLF-SOX families [31–39]. Recently, we have identified the 9aaTAD activation domain in MYCN and also in all Yamanaka transcription activators and showed the 9aaTADs activating transcription as short peptides [40].

In this study, we sought to investigate the activation of c-Myc transcription in constructs with and without the 9aaTAD. We discovered a strong interaction at nano-molar scale between the KIX domain of general coactivator CPB and the 9aaTAD activation domains of c-Myc and N-Myc proteins.

## Materials and Methods

### Expression construct and peptides

The GST-His6x-TEV-KIX construct was generated by PCR resulting in fusion product (GSTseq) - G S (linker) - H H H H H H (His6x tag) - G (linker) - E N L Y F Q / G (TEV tag) - (KIX region 586–672) and cloned into pGEX-2T vector. Custom peptide synthesis services were provided by Apeptide Shanghai Chutide Biotechnology Co., Ltd. Peptides have 95% purity and 2 mg were delivered #MLL: Biotin - GS-NILP-SDIM D FVLKNT, #c-Myc: Biotin - GSSS-TQLEMVTELLG, #N-Myc: Biotin - GSSS-EPP-SWVTEMLLEN, #SpQ: Biotin - GSS- GQ VSWQ T LQLQ NLQ, #SpD: Biotin - GSS-GD VSWD T LDLD DLD.

### Sample Preparation

The non-labelled KIX domain was expressed as described [38]. In brief, the uniformly labelled 13C, 15N labelled KIX domain used for assignment experiments and 15N labelled KIX domain used for titration experiments were expressed in M9 minimal media containing 15N-ammonium sulfate with and without 13C6-D-glucose, respectively. The cell pellet obtained from 1 L culture was re-suspended in 40 mL buffer (50 mM Tris-HCl, 150 mM NaCl, 10% glycerol, 40 mM imidazole, 3 mM NaN3, pH 7.5). Lysozyme was added to the pellet to the final concentration of 0.3 mg/mL and sonicated at 40 amplitude 1s on, 5s off while kept on ice. The lysate was centrifuged at 21,000 g for 1 hour at 4 °C. The supernatant was loaded on the His trap HP 5 mL column packed with Nickel (GE Healthcare). FPLC was used for elution of the GST-His6x-TEV-KIX protein with 300 mM imidazole in 50 mM Tris-HCl buffer (150 mM NaCl, 3 mM NaN3, pH 7.5). The GST-His6x-tag was removed by TEV protease overnight at 4 °C while dialyzing in 50 mM Tris-HCl buffer (150 mM NaCl, 3 mM NaN3, pH 7.5). Cleaved and dialyzed protein was loaded on the His trap HP 5 mL column packed with Nickel resin equilibrated with 50 mM Tris-HCl buffer (150 mM NaCl, 3 mM NaN3, pH 7.5) and the KIX domain was eluted with 150 mM imidazole in 50 mM Tris-HCl buffer (150 mM NaCl, 3 mM NaN3, pH 7.5). Eluted KIX domain was loaded on HiLoad S30 120 mL equilibrated with 20 mM NaPi buffer (50 mM NaCl, 1 mM NaN3, pH 6.0). The purity of protein samples between purification steps was verified by SDS PAGE and MALDI-TOF mass spectrometer and the quantity of KIX domain was determined by 280 nm absorbance. The quality of the protein was determined using Differential Scanning Fluorimetry (Prometheus NT.48, NanoTemper Technologies GmbH). The uniform labelling of KIX domain was confirmed by MALDI-TOF MS.

### Assessment of enzyme activities

The β-galactosidase activity was determined in the yeast strain L40 [34]. The strain L40 has integrated the lacZ reporter driven by the lexA operator. In all hybrid assays, we used 2μ vector pBTM116 for generation of the LexA hybrids. The yeast strain L40, the Saccharomyces cerevisiae Genotype: MATa ade2 his3 leu2 trp1 LYS::lexA-HIS3 URA3::lexA-LacZ, is deposited at ATCC (#MYA-3332). For β-galactosidase assays, overnight cultures propagated in YPD medium (1% yeast extract, 2% bactopeptone, 2% glucose) were diluted to an A600 of 0.3 and further cultivated for two hours and collected by centrifugation. The crude extracts were prepared by vortexing with glass beads for 3 minutes. The assay was done with 10 ul crude extract in 1ml of 100 mM phosphate buffer pH7 with 10 mM KCl, 1 mM MgSO4 and 0.2% 2-Mercaptoethanol; reaction was started by 200 ul 0.4% ONPG and stopped by 500 ul 1 M CaCO3. The average value of the β-galactosidase activities from two independent transformants is presented as a percentage of the reference with the standard deviation (means and plusmn; SD; n = 2). We standardized all results to previously reported Gal4 construct HaY including merely the activation domain 9aaTAD with the activity set to 100% [34].

### Bio-layer interferometry

Bio-layer interferometry on Octet RED96e (ForteBio) was used for the analysis of interaction between KIX domain and proposed activation domains. All experiments were performed at 30°C at 1000 rpm shaking. Streptavidin-bearing SA biosensors (ForteBio) were immersed for 300 sec into the assay buffer (20mM sodium phosphate pH 6.0, 50mM NaCl) containing 60 μM biotinylated MLL, MYC or NMY peptide, respectively. After subsequent wash in the assay buffer, the final steady response of ligand reached 0.6-0.8 nm. Parallel blank sensor was treated the same way with pure assay buffer being used instead of biotinylated peptide solution in the immobilization step. For the binding assay, all sensors were used in parallel applying the following procedure: 120 sec baseline in the assay buffer, 180 sec association in the assay buffer containing increasing concentration of KIX domain for each cycle (0.31, 0.63, 1.25, 2.5 and 5.0 μM), 240 sec dissociation in the assay buffer and 3 repetitions of 30 sec regeneration in 50mM NaOH followed by 30 sec assay buffer wash. Each binding experiment was performed in pentaplicate and the data were processed using Data Analysis 11.1 evaluation SW (ForteBio). Obtained binding curves were blank-subtracted and fitted by a 1:1 binding model using steady state analysis. Final KD (apparent) values were calculated as an average of all measurements.

### Western Blot Analysis

The desaturated yeast total protein samples were prepared by heating cells in lysis buffer at 94°C for 5 minutes (lysis buffer: 10% SDS, 500 mM Tris-HCl pH 6.8, 500 mM DTT, 50% glycerol v/v, 0.025% bromophenol blue dye). Proteins from samples were spread out by molecular size during SDS-PAGE and blotted to nitrocellulose. The immuno-detection of proteins was carried out using mouse anti-HA monoclonal antibody (2-2.2.14, #26183, ThermoFisher Sci) and secondary anti-mouse IgG antibodies conjugated with horseradish peroxidase (#A9044, Sigma Aldrich). The proteins were visualized using Pierce ECL (#32106, ThermoFisher Sci) according to the manufacturer’s instructions. Noteworthy, we observed higher abundance of c-Myc protein generated from both shorter constructs (69-103 and 69-108), which expressions were significantly higher than the longer c-Myc constructs (1-103 and 1-108).

## Results

### The 9aaTAD is well conserved in evolution of the MYC family

Previously, we have identified the 9aaTADs in human c-Myc and N-Myc and also in all three Myc animal paralogs in lobe-finned bony fish coelacanth (including functional L-Myc) and experimentally determined the 9aaTADs activating transcription as short peptides [37]. In higher metazoans, including humans, the 9aaTAD activation domains is completely absent in L-Myc paralogs (Fig 1a), what suggests that the loss of 9aaTAD in L-Myc paralogs has occurred already in early tetrapod evolution. In lower metazoans, including all fishes, we found the conservation of the 9aaTAD in all members of the Myc family and that to the last unicellular ancestor of animals, choanoflagellates, represented here by Monosiga brevicollis (Fig 2).

**Figure 1.**
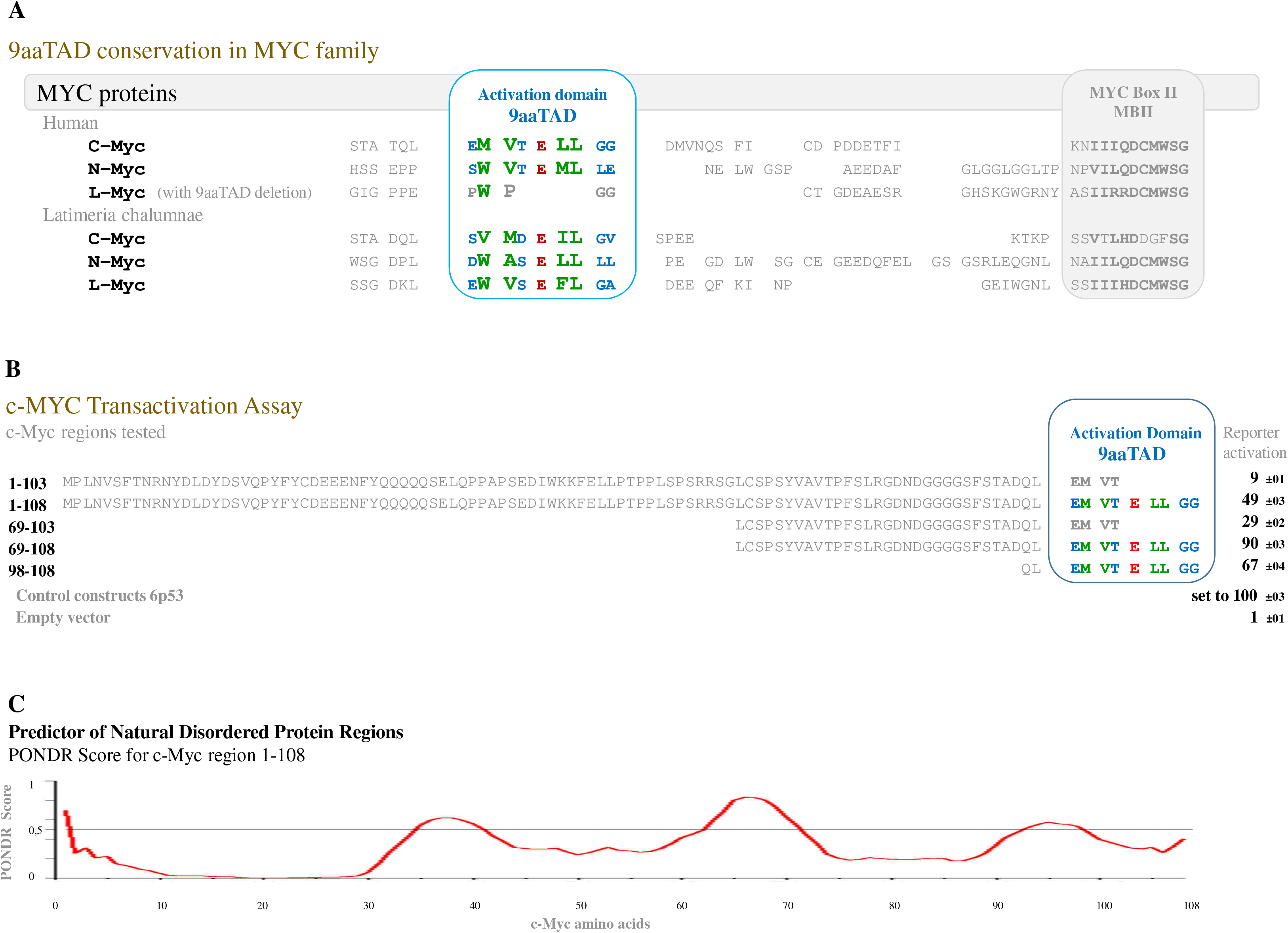
Conservation of the 9aaTAD in MYC family. **A**, Conservation of the 9aaTAD domain in MYC family. We found conservation of the activation domains in all three Myc paralogs in lobe-finned bony fish coelacanth (Latimeria chalumnae, fin-to-limb transition, related to lungfishes and tetrapods), which also conserved in human c-Myc and N-Myc but lost in L-Myc. The 9aaTADs activation domains are colored for faster orientation. The 9aaTAD deletion in L-Myc is in grey. **B**, Transactivation Assay. The activation of transcription by identified 9aaTAD alone (region 98-108) is comparable with entire N-terminal region of c-Myc (region 98-108). The regions with the c-Myc 9aaTAD activation domains were tested in a reporter assay with hybrid LexA DNA binding domain for the capacity to activate transcription. The average value of the β-galactosidase activities from two independent transformants is presented as a percentage of the reference with standard deviation (means and plusmn; SD; n = 3). We standardized the results to positive control p53 construct 6p53, which was set to 100% (raw value 273 +/-9) and control empty vector Hdd (raw value 1,5 +/-1). The 9aaTADs activation domains are colored for faster orientation. Deuterated 9aaTADs or their partial deletion are in grey. **C**, Predictor of Natural Disordered Protein Regions (PONDR). Prediction result for c-myc region 1-108 is shown. PONDR® is copyright ©1999 by the WSU Research Foundation.

**Figure 2.**
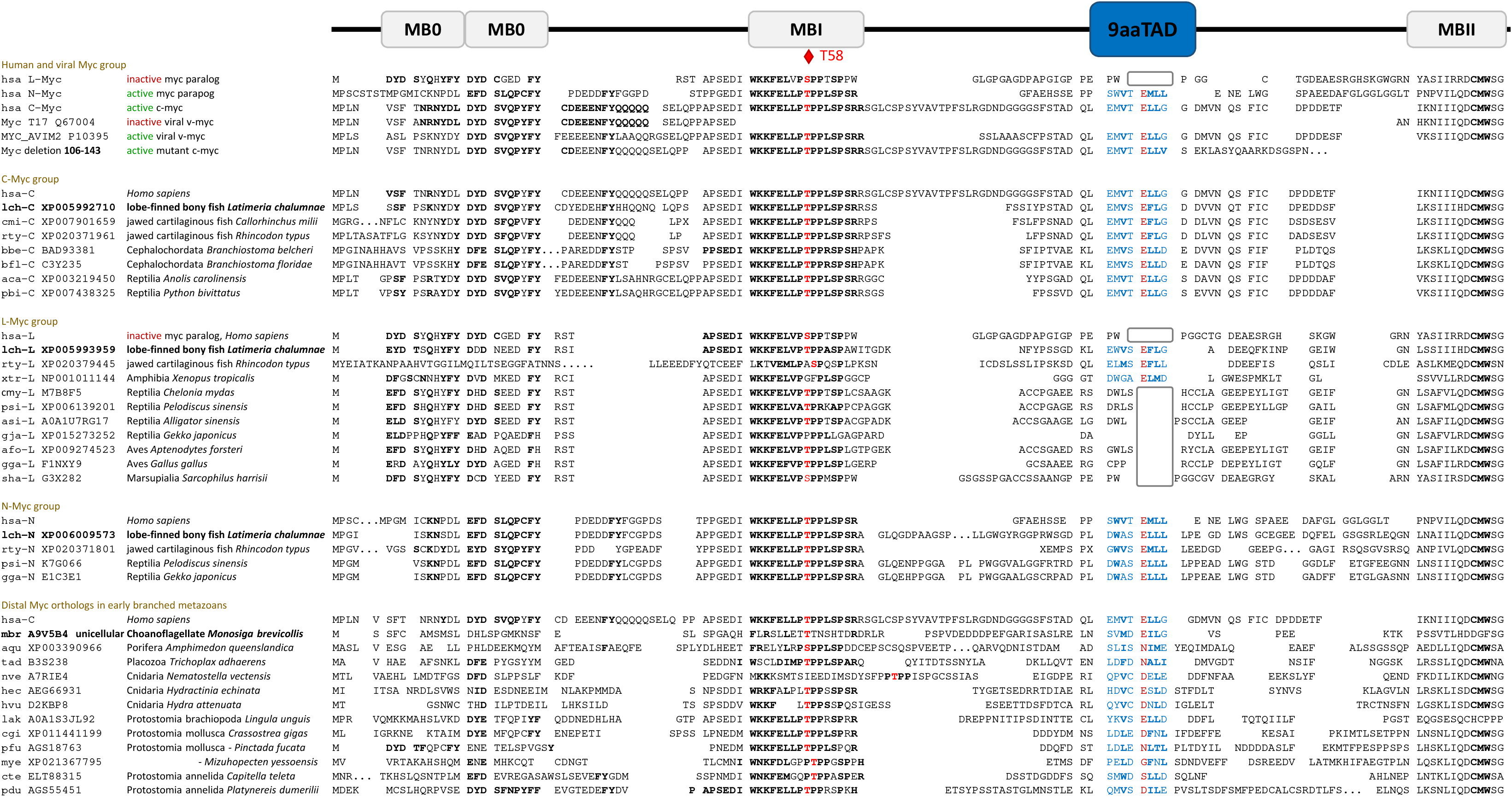
Alignment of the MYC family. The N-terminal regions of MYC proteins were aligned by sequence similarities and their predicted 9aaTAD activation domains are shown. The 9aaTADs activation domains are colored for faster orientation. The MYC clades with conservation MYB box O, I and II are shown. The deletion of activation domain in L clade are in grey box. Conservation of threonine in position 55 is highlighted in red.

### Activation of c-Myc transcription is linked with the 9aaTAD

Next, we tested the N-terminal regions of c-Myc as activators of transcription. The alone-standing c-Myc 9aaTAD activation domain (region 98-108), the c-Myc N-terminal regions with 9aaTAD (1-108, 69-108) and corresponding c-Myc regions without 9aaTAD (1-103, 69-103). All generated constructs were tested for activation of transcription in one hybrid assay (Fig. 1b). The constructs without 9aaTAD have lost the majority of their transcriptional capacity in comparison with corresponding constructs including the 9aaTAD. Furthermore, we observed higher abundance of c-Myc protein generated from both shorter constructs (69-103 and 69-108), which expressions were significantly higher than the longer c-Myc constructs (1-103 and 1-108) and their expression were below our immuno-detection threshold (Supplementary Figure S1). The naturally disordered prediction for c-Myc region 1-108 was generated by PONDR algorithm [41] (Fig. 1c).

### The KIX domain interactions with c-Myc and N-Myc

Next, we sought to investigate the KIX interactions with the 9aaTADs by bio-layer interferometry (BLI). As MLL and other members of the 9aaTAD family bind to KIX domain of CBP [42], we used well-studied MLL peptide as a positive control here and previously [38]. The MLL 9aaTADs occupied the same space on the KIX domain as E2A and p53. Furthermore, they induced the KIX intramolecular reformation realized by the two-point interaction involving 9aaTAD positions p3-4 and p6-7 [38]. For all three tested 9aaTADs, a clear binding to the KIX was observed (Fig. 3a).

**Figure 3.**
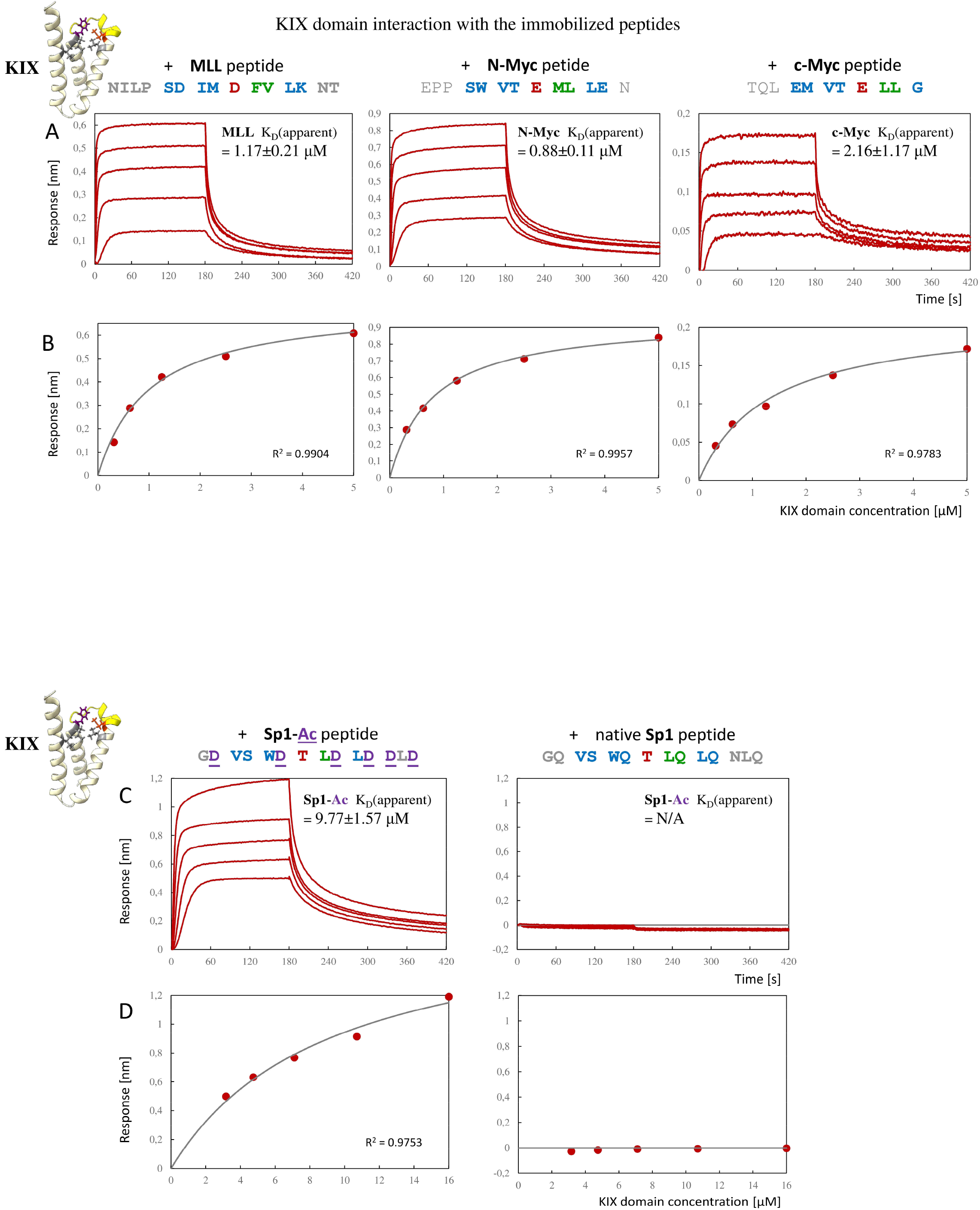
Bio-layer interferometry assay (BLI). A single experiment of pentaplicate is shown for each of the tested peptides (MLL, N-Myc, c-Myc, Sp1-Ac, Sp1). **A** and **C**, blank-subtracted association (180 sec) and dissociation (240 sec) curves for 5 concentrations of KIX domain interaction with the immobilized peptide. **B** and **D**, steady-state analysis of corresponding measurement in panel A. The response for each concentration shown as red circle, fitted curve for 1:1 binding model shown in grey. The goodness of each curve fit is given by the coefficient of determination R2. Values close to 1.0 indicate proximity to 100% fit.

Through steady-state analysis of the BLI-binding assay, the equilibrium dissociation constant (Kd) values in the micromolar to sub-micromolar range were discovered. The N-Myc peptide displayed Kd(apparent) of 0.88±0.11 μM and c-Myc peptide Kd(apparent)= 2.16±1.17 μM. The BLI measurements for immobilized MLL peptide and KIX protein (Kd(apparent)= 1.17±0.21 μM) are in good agreement with our prior reference’s measurement using isothermal titration calorimetry for free MLL peptide and KIX protein (Kd= 1.4±0.1 μM) [38]. Final KD (apparent) values were calculated as an average of all measurements (for more details see Methods).

In conclusion, both Myc peptides displayed strong binding to the KIX comparable to MLL peptide. Each binding experiment was performed in pentaplicate and the data were processed using Data Analysis 11.1 evaluation SW-ForteBio. Obtained binding curves were blank-subtracted (including Streptavidin-bearing SA-biosensors and KIX but excluding peptide) and fitted by a 1:1 binding model using steady state analysis (Fig. 3b).

### Glutamines prevent Sp1 from KIX binding

To further confirm the above results and demonstrate the unbiased BLI-assay outcomes, we carried out additional experiment with activation domain that does not bind to KIX. Previously, we addressed the importance of over-represented glutamines in Sp1 activation region for ability to activated transcription (37). In this study, we substituted the over-represented glutamines for aspartic acid residues (construct Sp1-Ac, acidic form of Sp1 9aaTAD) (Fig. 3c).

We tested native Sp1-9aaTAD and the acidic modification for KIX binding by BLI-assay. The result from BLI-measurement for Sp1-Ac peptide and KIX (Kd(apparent)= 9.77±1.57 μM) was comparable to MLL (Kd(apparent)= 1.17±0.21 μM) but the native Sp1 did not bind to KIX in the 3-16 μM range (Kd not determined) and served as the negative control for all BLI-measurements (Fig. 3c, d). These results confirmed that the overabundance of glutamine residues prevents binding to the KIX domain.

## Discussion

The ongoing efforts to identify molecules able to directly bind, interfere and/or suppress MYC activity have been the scope of intense research in numerous laboratories. The Omomyc, a 90-residue peptide, was shown to bind Myc by interfering with dimerization with its partner MAX and thus suppressing cell proliferation [24].

Besides the attempts to target Myc directly, another approach was to suppress MYC activity by targeting factors regulating MYC degradation [43], alternative strategies target MYC stability or blocking its interaction between acetylated histones and the bromodomain protein BRD4 [44]. The JQ1, a small molecule inhibitor of BET proteins, has been shown to suppress Myc expression and cancer cell growth in vitro and in vivo in preclinical models of multiple myeloma, AML and CML [26–28]. Ongoing studies among others led by Dr Laura Soucek [45] and future studies including the focus on the 9aaTADs c-Myc and the KIX interaction will aim to translate new findings into development of small drug inhibitors and can guide therapies targeting Myc-dependent tumors.

### Conclusions

In this study, we tested the human c-Myc constructs (1-108, 69-108 and 98-108) in N-terminal regions for their ability to activate transcription. We discovered a very strong interaction at nano-molar scale between the KIX domain of general coactivator CPB and the 9aaTADs of c-Myc and N-Myc proteins. The close-fitting Myc and KIX binary interaction represents new promising druggable target, which could initiate further studies.

## Abbreviations

9aaTAD: nine-amino-acid activation domain
AML: acute myeloid leukemia
BLI: bio-layer interferometry
CML: chronic myeloid leukemia
MB: conserved regions in Myc proteins, Myc boxs
Kd: equilibrium dissociation constant
KIX: binding domain in CBP a P300 proteins

## Declarations

### Ethics approval and consent to participate

Not relevant for this study!

### Consent for publication

Not applicable.

### Availability of data and materials

The datasets used and/or analyzed during the current study are available from the corresponding author on reasonable request.

### Author’s Contributions

MP and AK conceived and designed the project. JH and MP performed the experiments. VJ, VV and TO discussed the data. JH, MP and AK wrote the manuscript. All authors have contributed critically to intellectual content and have approved the final manuscript.

## Acknowledgements

We thank Alena Hofrova for technical support and Apeptide Shanghai Chutide Biotechnology for excellent quality of supplied peptides.

