## Supplementary Figure S1 for "Undruggable oncoproteins cMyc and NMyc bind to mediator of transcription with superior high affinity"

### Suppl. Figure S1

**Westernblotting:**
(anti-HA antibody)

250 kDa
130 kDa
100 kDa
70 kDa
55 kDa
35 kDa
25 kDa
15 kDa

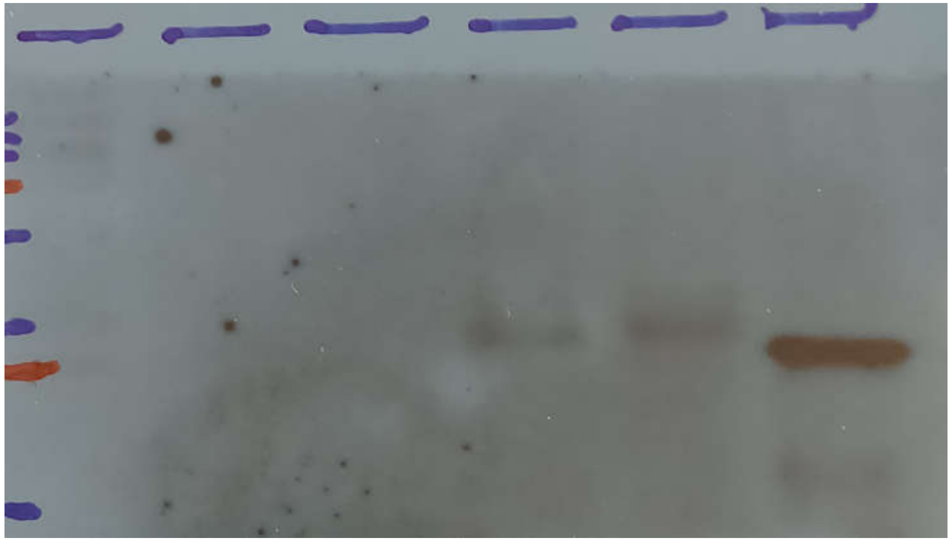

LexA-HA-cMyc
LexA-HA

| Constructs LexA - HA- cMyc regions: | (Marker) | (1-103) | (1-108) | (69-103) | (69-108) | (0-0) |
|---|---|---|---|---|---|---|
| Expected size in kDa | | 36,7 | 37,1 | 28,6 | 29,0 | 25,7 |

**Westernblotting - repeat:**
(anti-HA antibody)

250 kDa
130 kDa
100 kDa
70 kDa
55 kDa
35 kDa
25 kDa
15 kDa

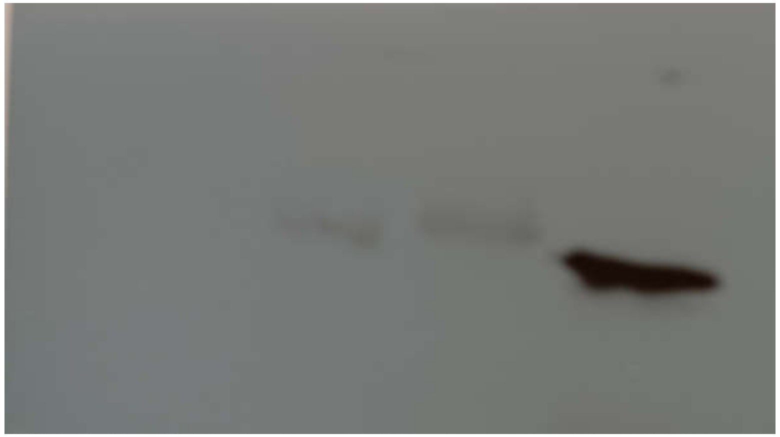

LexA-HA-cMyc
LexA-HA
